# BubR1 TPR domain supports mitotic checkpoint by promoting MCC formation and MCC-APC/C interaction

**DOI:** 10.1101/2025.10.13.682252

**Authors:** Mingzhe Zhang, Tingting Lei, Ying Wang, Mingyan Li, Yutong Wang, Jing Fang, Gang Zhang

## Abstract

BubR1 is the key component of the mitotic checkpoint, a surveillance mechanism ensures accurate chromosome segregation by facilitating the assembly of the mitotic checkpoint complex and promoting its binding to the anaphase-promoting complex/cyclosome. Although BubR1’s role in SAC signaling has been extensively investigated, the function of its N-terminal tetratricopeptide repeat (TPR) domain remains poorly understood.

In this study, we first established the essential role of the BubR1 TPR domain in SAC signaling. Guided by the resolved cryo-EM structure of the MCC-APC/C complex, we identified and characterized several interactions involving this domain with Mad2, Cdc20^APC/C^, Apc2. Furthermore, we discovered an intramolecular interaction between the TPR domain and downstream residues of BubR1, which appears to organize a structure resembling a “lasso” that incorporates four Cdc20^APC/C^-binding elements, thereby enhancing engagement with Cdc20 in the APC/C. Functional and biochemical analyses demonstrated that these interactions collectively promote MCC assembly and MCC-APC/C binding, enabling rapid SAC activation in response to microtubule-kinetochore attachment defects.

## Introduction

To maintain genome stability, eukaryotic cells have evolved the spindle assembly checkpoint (SAC), a signaling pathway that minimizes chromosome segregation errors during cell division. In the presence of unattached kinetochores, this pathway is activated leading to the formation of the mitotic checkpoint complex (MCC), which binds and inhibits the anaphase promoting complex/cyclosome (APC/C), an E3 ligase required for ubiquitination of securin and cyclin B1. By preventing the degradation of these key mitotic regulators, the SAC maintains cells in mitosis until all kinetochores are properly attached to spindle microtubules (Musacchio, 2015).

After three decades of extensive investigation, the molecular mechanism of SAC activation has been progressively elucidated. Through the action of mitotic kinases such as Mps1 and Cdk1, key checkpoint proteins including Bub1, BubR1, Mad1, Mad2, and Cdc20 are sequentially recruited to unattached outer kinetochores, primarily via phosphorylation-dependent interactions (Saurin, 2018). At the kinetochores, Bub1 acts as a scaffold to bring Mad1/Mad2 molecules close to Cdc20, facilitating the formation of Mad2-Cdc20 complex (Ji et al., 2017; Zhang et al., 2017; Faesen et al., 2017; Lara-Gonzalez et al., 2021; Piano et al., 2021; Fischer et al., 2021; Fischer et al., 2022; Chen et al., 2023; Sethi et al., 2025; Yu et al., 2025). Afterwards, the Mad2-Cdc20 complex associates with BubR1-Bub3 to form the mature MCC (Kulukian et al., 2009; Faesen et al., 2017). BubR1, a key component of the MCC, is largely intrinsically disordered except for a structured N-terminal TPR (tetratricopeptide repeat) domain (residues 59-219) and a C-terminal pseudokinase domain. Within its flexible region, BubR1 harbors multiple short linear interaction motifs (SLiMs) that are essential for SAC signaling and chromosome alignment. These include two KEN boxes (KEN1: 26-28; KEN2: 304-306), two D boxes (D1: 224-232; D2: 555-563), and three ABBA motifs (ABBA1: 272-277; ABBA2: 340-345; ABBA3: 528-533), which collectively engage two molecules of Cdc20 to facilitate MCC assembly and APC/C inhibition (Burton et al., 2007; King et al., 2007; Sczaniecka et al., 2008; Malureanu et al., 2009; Elowe et al., 2010; Lara-Gonzalez et al., 2011; Lischetti et al., 2014; Di Fiore et al., 2015; Diaz-Martinez et al., 2015; Izawa and Pines 2015; Di Fiore et al., 2016; Alfieri et al., 2016; Sewart et al., 2017). Additionally, the BubR1 GLEBS motif (residues 408-418) interacts with Bub3 and, together with the Bub1-binding motif (residues 440-460), governs kinetochore targeting (Taylor et al., 1998; Wang et al., 2001; Larsen et al., 2007; Zhang et al., 2016). An LxxIxE motif (also termed the KARD motif, residues 669-674) mediates direct binding to PP2A-B56, which promotes kinetochore-microtubule attachment and SAC silencing (Suijkerbuijk et al., 2012; Kruse et al., 2013; Xu et al., 2013).

In addition to these well-characterized SLiMs, several unresolved questions persist regarding the N-terminal region of BubR1. Whether the TPR domain is required for SAC signaling is unknown yet. Early studies identified a direct interaction between the BubR1 TPR domain and the KNL1 KI2 motif (residues 238-249) and proposed that this interaction might be important for BubR1 kinetochore localization and SAC signaling (Kiyomitsu et al., 2007; Kiyomitsu et al., 2011; Bolanos-Garcia et al., 2011; Krenn et al., 2012). However, subsequent work revealed that phosphorylated MELT motifs in KNL1 serve as the primary docking sites for Bub3, which is critical for BubR1 recruitment to kinetochores (London et al., 2012; Shepperd, et al., 2012;Yamagishi et al., 2012). Functional analyses further indicated that the KI motifs contribute only marginally to SAC signaling in the context of full length KNL1 (Vleugel et al., 2013) or exhibit minor effects in truncated N-terminal fragment (Krenn et al., 2014). Whether the corresponding residues in BubR1 TPR domain participate in SAC signaling and the underlying mechanism remain uncharacterized. Another unresolved issue concerns the direct interaction between the BubR1 N-terminal region and Mad2. Although an *in vitro* interaction between BubR1^1-371^ and Mad2 has been reported (Tipton et al., 2011), its *in vivo* relevance, structural basis, and functional significance remain unclear. Additionally, the highly conserved residue W295 in BubR1 has been shown to be essential for SAC signaling (Tromer et al., 2016), yet the underlying mechanism remains unknown. In a related context, a missense mutation of R286S in Cdc20 was identified in a patient with premature aging syndrome (Fujita et al., 2020). While the authors attributed its effect to disrupted interaction with K26 in the BubR1 KEN1 box, this assignment conflicts with structural analyses (Tian et al., 2012; Alfieri et al., 2016;2020). Therefore, a re-examination of the interaction between Cdc20 R286 and BubR1 is warranted.

The cryo-EM structure of MCC-APC/C complex has been resolved at near-atomic resolution (Alfieri et al., 2016; 2020). Although the middle and C-terminal regions of BubR1 remain unresolved in this structure, its entire TPR domain of BubR1 is well defined, providing a structural roadmap to address the outstanding questions outlined above. In this study, we first established the functional importance of the TPR domain itself, demonstrating that it is essential for SAC signaling. We then systematically characterized a set of interactions involving the BubR1 TPR domain with Mad2, Cdc20^APC/C^, and Apc2, as inferred from the cryo-EM map. Our functional analyses confirm that these interactions are critical for SAC and reveal their underlying mechanisms. Notably, we discovered an intramolecular interaction between the TPR domain and several residues preceeding KEN2 box, including W295, which together form a structural element resembling a “lasso”. This lasso structure contains all the three degrons (A1, D2 and K2) involved in Cdc20^APC/C^ binding, as well as residues E134 and D137, which form an electrostatic interface with Cdc20^APC/C^ R286. This configuration likely enhances the efficiency of Cdc20^APC/C^ engagement, enabling a rapid checkpoint response as shown by mutation analysis. In summary, our work elucidates essential roles and detailed mechanisms of the BubR1 TPR domain in promoting both MCC assembly and MCC-APC/C interaction.

## Results

### The BubR1 TPR domain plays a critical role in SAC signaling

Mitotic checkpoint protein BubR1 contains an N-terminal TPR domain whose function in SAC signaling remains poorly understood (Fig. 1A). To investigate whether this domain is required for SAC signaling, we generated two BubR1 TPR mutants: BubR1ΔTPR lacking the entire TPR domain (Δ59-219) and BubR1 B1TPR, where the TPR domain was replaced by the structurally similar but sequence-divergent TPR domain from Bub1 (residues 1-150) (Fig. 1B). Both mutant proteins localized correctly to kinetochores (Supp. Fig. 1). To address their functional competence, we conducted RNAi and rescue experiment as described previously (Song et al., 2024). HeLa cells were co-transfected with BubR1-targeting siRNA and an RNAi-resistant, YFP-tagged BubR1 expression plasmid. Live cell imaging was used to measure the duration of mitotic arrest induced by a low dose of nocodazole (40 ng/ml), which depolymerizes spindle microtubules and activates the SAC. Mitotic arrest time serves as a direct indicator of checkpoint strength. Control cells exhibited a robust SAC, with a median mitotic arrest time of approximately 715 min, whereas BubR1-depleted cells completely lost checkpoint function, exiting mitosis within 20 min (Fig. 1C). Expression of wild-type BubR1 fully restored checkpoint activity, prolonging mitotic arrest to 660 min. In contrast, neither TPR mutant supported SAC function, despite their proper kinetochore localization (Fig. 1C). The failure of SAC activation could result from defects in MCC assembly or impaired MCC-APC/C binding. To explore the underlying cause for the SAC deficiency, we performed YFP-affinity purification of YFP-BubR1 and YFP-BubR1 B1TPR mutant, which more closely resembles full length BubR1 than the BubR1ΔTPR mutant from mitotic cells. We found that the BubR1 B1TPR mutant failed to form the MCC (Fig. 1D). This result suggests that the TPR domain is essential for MCC assembly, though it does not rule out an additional role in mediating MCC-APC/C interaction, which we further investigated below.

**Figure 1.**
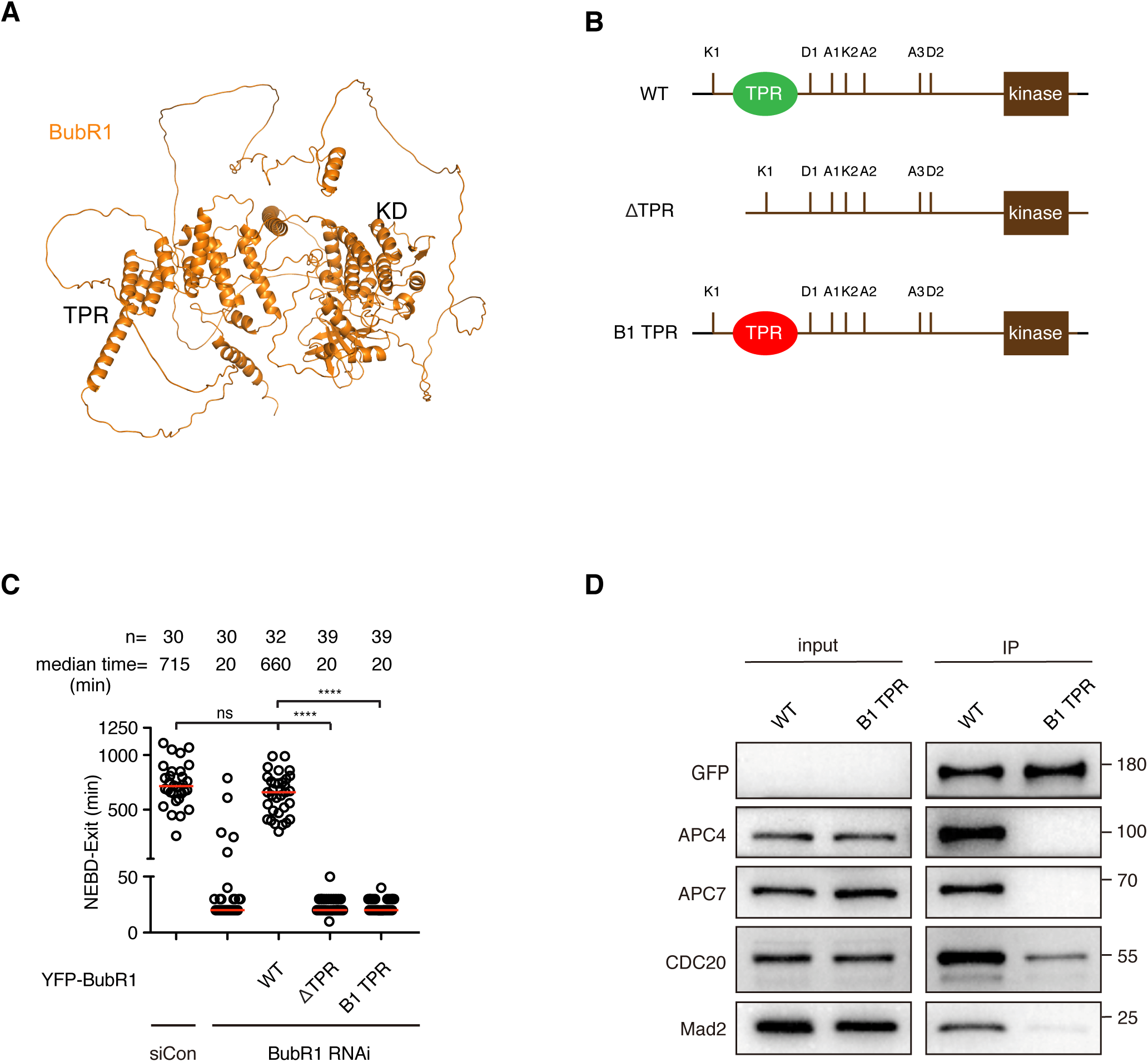
The BubR1 TPR domain is critical for SAC signaling. **A** Predicted BubR1 structure by AlphaFold. **B** Schemes showing structures of wild type BubR1 and two BubR1 TPR mutants. **C** Plot showing the time from NEBD to mitotic exit of the cells transfected with siRNA oligos against BubR1 and RNAi-resistant constructs expressing YFP-BubR1. Low dose of nocodazole (40 ng/ml) was applied into the medium before live cell imaging conducted. Each circle represents the time spent in mitosis of a single cell. Red line indicates the median time. The number of cells analyzed per condition is indicated above (n = X). Mann-Whitney u-test was applied. ns means not significant; **** means P<0.0001. **D** YFP-BubR1 or YFP-BubR1 B1TPR was expressed in HeLa cells and immunoprecipitated by GFP-trap beads. YFP-BubR1, APC4, APC7, Cdc20 and Mad2 were detected by western blot from the immunoprecipitate.

The above results clearly demonstrated an essential role for the BubR1 TPR domain in SAC signaling.

### The BubR1-Mad2 interaction is essential for SAC signaling

To further elucidate the molecular mechanism by which the TPR domain contributes to SAC signaling, we analyzed the reported structure of MCC-APC/C complex in details (Alfieri et al., 2016;2020). The structure reveals a helix-loop-helix (HLH) motif (residues 20-47) extending from the TPR domain. Within this HLH motif, the KEN1 box (residues 26-28) contacts with the KEN receptor surface on Cdc20^MCC^. Notably, two residues in the same helix, W22 and E23, are positioned in close proximity to F141 and R133 of Mad2, suggesting potential hydrophobic and electrostatic interactions respectively (Fig. 2A). To address whether this putative interaction is required for SAC signaling, we constructed two BubR1 mutants, E23A and W22A/E23A. Immunofluorescence analysis confirmed that both mutants localized properly to kinetochores (Supp. Fig. 1). However, functional analysis by live cell imaging revealed defective SAC signaling in both mutants (Fig. 2B). Immunoprecipitation of YFP-BubR1 and YFP-BubR1 W22A/E23A from mitotic cells showed that the W22A/E23A mutant failed to form the MCC, explaining the loss of checkpoint activity (Fig. 2C). To determine whether these residues mediate direct interaction between BubR1 and Mad2, we purified recombinant His-tagged Mad2 L13A, a mutant that keeps Mad2 in the closed conformation and GST-tagged wild type or W22A/E23A BubR1 (residues 1-220), and performed GST pull-down assay. The result showed a pronounced reduction in Mad2 binding to the W22A/E23A mutant compared to the wild type protein (Fig. 2D).

**Figure 2.**
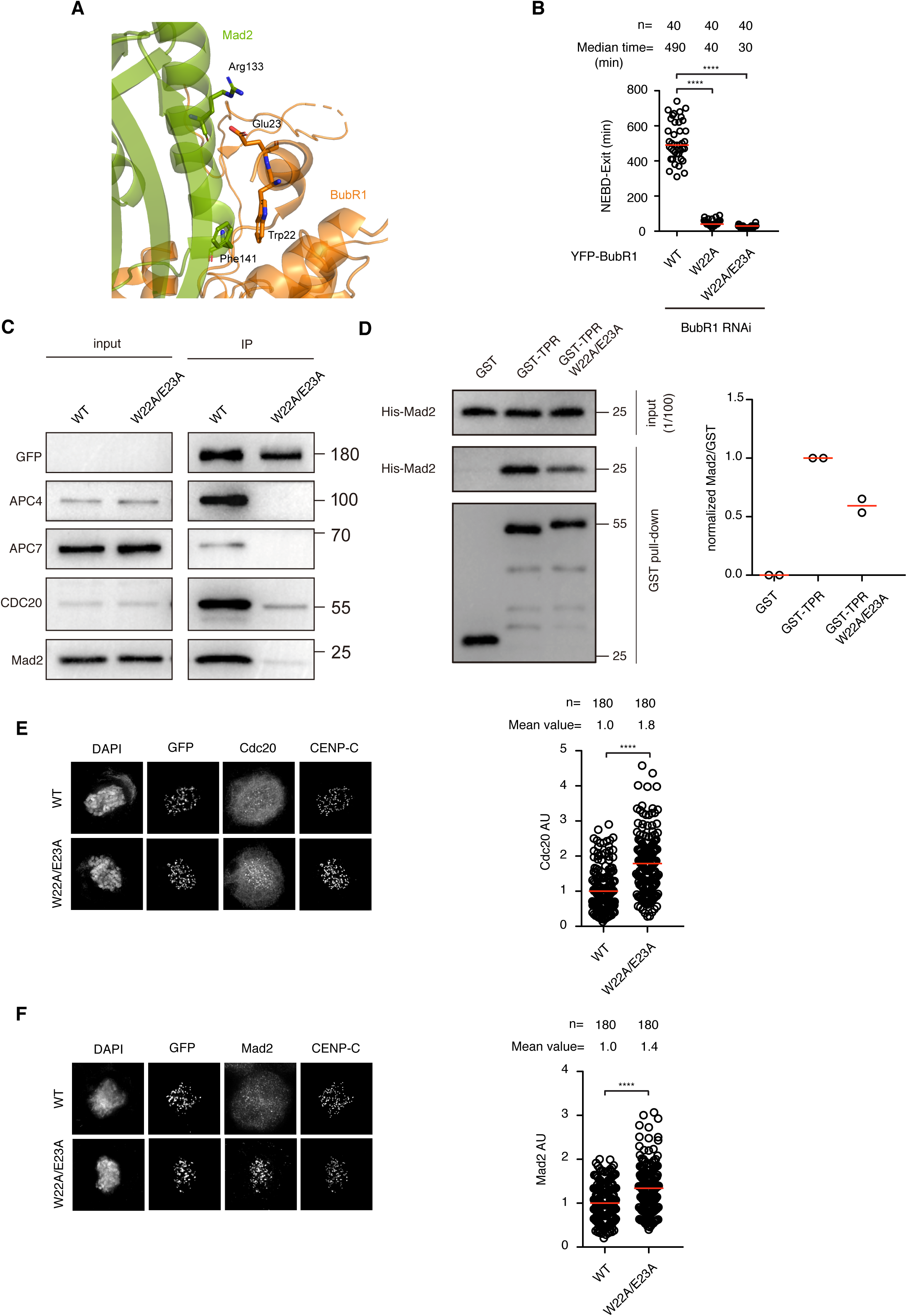
The BubR1 TPR-Mad2 interaction plays essential role in SAC signaling. **A** Structural analysis of BubR1 TPR-Mad2 interaction from PDB: 6TLJ. **B** Plot showing the time from NEBD to mitotic exit of the cells transfected with siRNA oligos against BubR1 and RNAi-resistant constructs expressing YFP-BubR1 E23A or W22A/E23A. Low dose of nocodazole (40 ng/ml) was applied into the medium before live cell imaging conducted. Each circle represents the time spent in mitosis of a single cell. Red line indicates the median time. The number of cells analyzed per condition is indicated above (n = X). Mann-Whitney u-test was applied. **** means P<0.0001. **C** YFP-BubR1 or YFP-BubR1 W22A/E23A was expressed in HeLa cells and immunoprecipitated by GFP-trap beads. YFP-BubR1, APC4, APC7, Cdc20 and Mad2 were detected by western blot from the immunoprecipitate. **D** *in vitro* protein binding assay showing direct interaction between BubR1 TPR and Mad2. Left: western blot showing the presence of His-tagged Mad2 pulled down together with GST-tagged BubR1 TPR recombinant protein. Right: quantification of the western blot results from two repeats. The line represents the median value of two repeats. The value of GST alone was substracted from the values of the GST-BubR1 TPR recombinant proteins and set to zero. The value from wild type GST-BubR1 TPR was set to 1 and the value from GST-BubR1 TPR 2A was normalized to the wild type value. **E** Immunofluorescence of kinetochore signals of Cdc20 in HeLa cells depleted of endogenous BubR1 by RNAi and supplemented with exogenous YFP-BubR1. Left: representative images of the cells stained with individual antibodies. Right: quantification of kinetochore signals of Cdc20. 180 kinetochores from 12 cells were quantified and plotted in each condition. **F** Immunofluorescence of kinetochore signals of Mad2 in HeLa cells treated similarly as in **E**. Left: representative images of the cells stained with individual antibodies. Right: quantification of kinetochore signals of Cdc20. 180 kinetochores from 12 cells were quantified and plotted in each condition.

We next asked whether this interaction occurs at kinetochores. To test this, we depeleted endogenous BubR1 and expressed either YFP-BubR1 or YFP-BubR1 W22A/E23A, and then examined kinetochore levels of Mad2 and Cdc20. Indeed, cells expressing the BubR1 W22A/E23A mutant exhibited significantly elevated kinetochore signals for both Mad2 and Cdc20 (Fig. 2E,F).

### The BubR1 TPR-Apc2 interaction is important for SAC signaling

Structural analysis also suggested a potential interaction between the BubR1 TPR domain and the APC/C component Apc2. At least four residues in the BubR1 TPR domain, R169, F175, V200 and L205 appear to participate in the interaction (Fig. 3A). Mutating all the four residues (4A) did not disrupt kinetochore localization but completely abrogated SAC signaling (Supp. Fig. 1; Fig. 3B). Surprisingly, immunoprecipitation assays revealed that the 4A mutation not only disrupted APC/C binding but also severely impaired MCC assembly (Fig. 3C). These pleiotropic effects made it difficult to specifically assess the contribution of the BubR1-Apc2 interaction on SAC signaling using this mutant. To uncouple MCC formation from APC/C binding, we generated more refined mutations: R169A/F175A and V200A/L205A. Live cell imaging showed that SAC activity was strongly impaired in the R169A/F175A mutant (median mitotic arrest around 70 min vs 515 min for wild type), but only mildly reduced in the V200A/L205A mutant (around 185 min) (Fig. 3D). Immunoprecipitation analysis confirmed that both mutants significantly reduced APC/C binding. However, while MCC formation remained intact in the V200A/L205A mutant, it was disrupted in the R169A/F175A mutant (Fig. 3E). These results indicate that the BubR1 TPR-Apc2 interaction indeed plays an important role in SAC signaling, likely by promoting the MCC-APC/C interaction.

**Figure 3.**
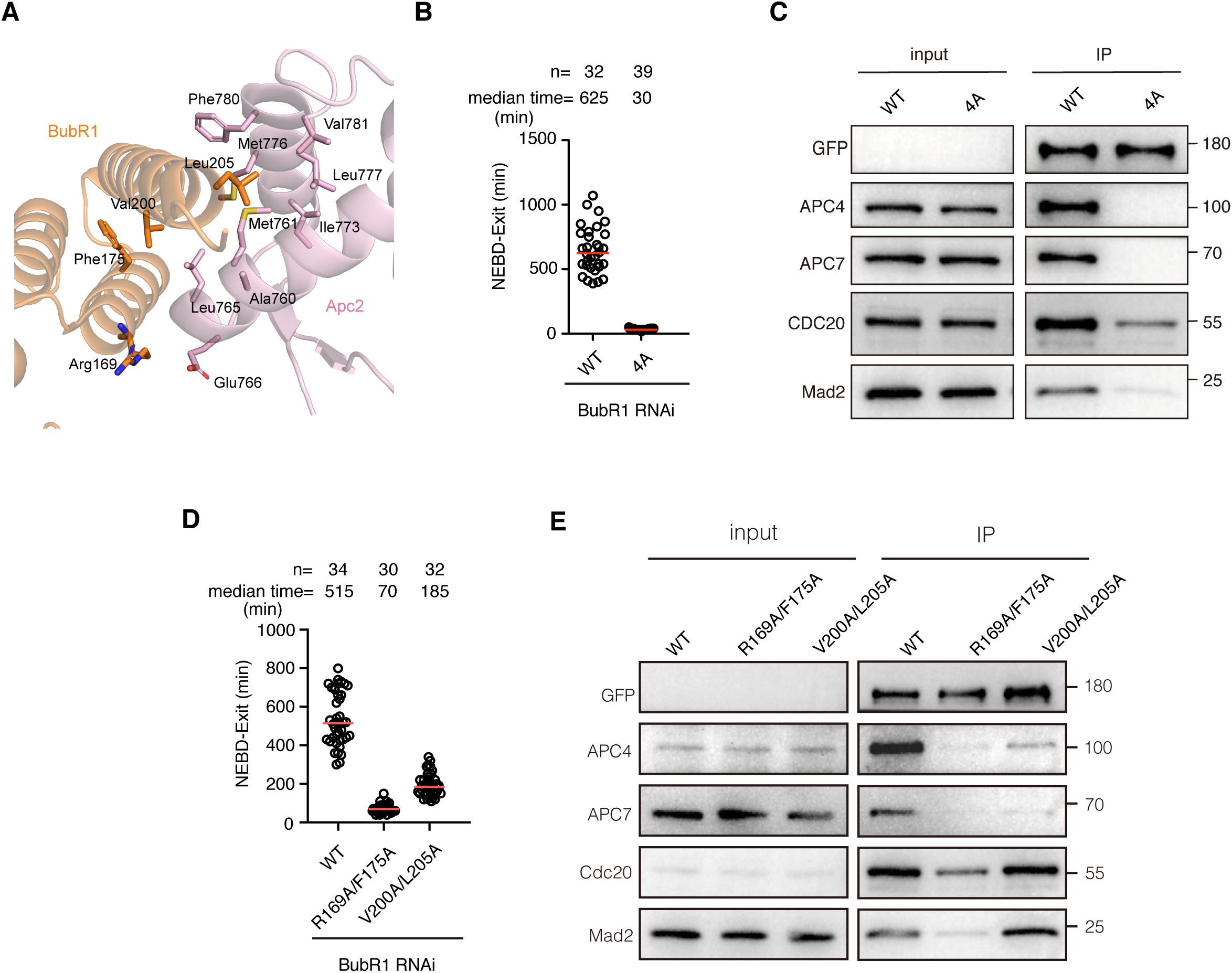
The BubR1 TPR-Apc2 interaction performs important role in SAC signaling. **A** Structural analysis of BubR1 TPR-Apc2 interaction from PDB: 6TLJ. **B** Plot showing the time from NEBD to mitotic exit of the cells transfected with siRNA oligos against BubR1 and RNAi-resistant constructs expressing YFP-BubR1 or YFP-BubR1 R169A/F175A/V200A/L205A (4A). Low dose of nocodazole (40 ng/ml) was applied into the medium before live cell imaging conducted. Each circle represents the time spent in mitosis of a single cell. Red line indicates the median time. The number of cells analyzed per condition is indicated above (n = X). Mann-Whitney u-test was applied. **** means P<0.0001. **C** YFP-BubR1 or YFP-BubR1 4A was expressed in HeLa cells and immunoprecipitated by GFP-trap beads. YFP-BubR1, APC4, APC7, Cdc20 and Mad2 were detected by western blot from the immunoprecipitate. **D** Plot showing the time from NEBD to mitotic exit of the cells treated similarly as in **B**. **E** Immunoprecipitation of YFP-BubR1 or YFP-R169A/F175A or YFP-BubR1 V200A/L205A in cells treated similarly as in **C**.

### The BubR1 TPR-Cdc20^APC/C^ interaction is critical for SAC signaling

Structural analysis further identified a potential electrostatic interaction between BubR1 TPR residues E134, D137 and residue R286 of Cdc20^APC/C^ (Fig. 4A). Disrupting this interaction by substituting E134 and D137 in BubR1 with positively charged residues had no effect on kinetochore localization, but completely abolished SAC signaling, a phenotype similarly observed when replacing Cdc20 R286 with a negatively charged residue (Supp. Fig 1; Fig. 4B,C). Consistent with structural and functional data, immunoprecipitation assay showed that the BubR1 E134R/D137R mutant retained the ability to form the MCC, albeit less efficiently, but failed to bind the APC/C (Fig. 4F).

**Figure 4.**
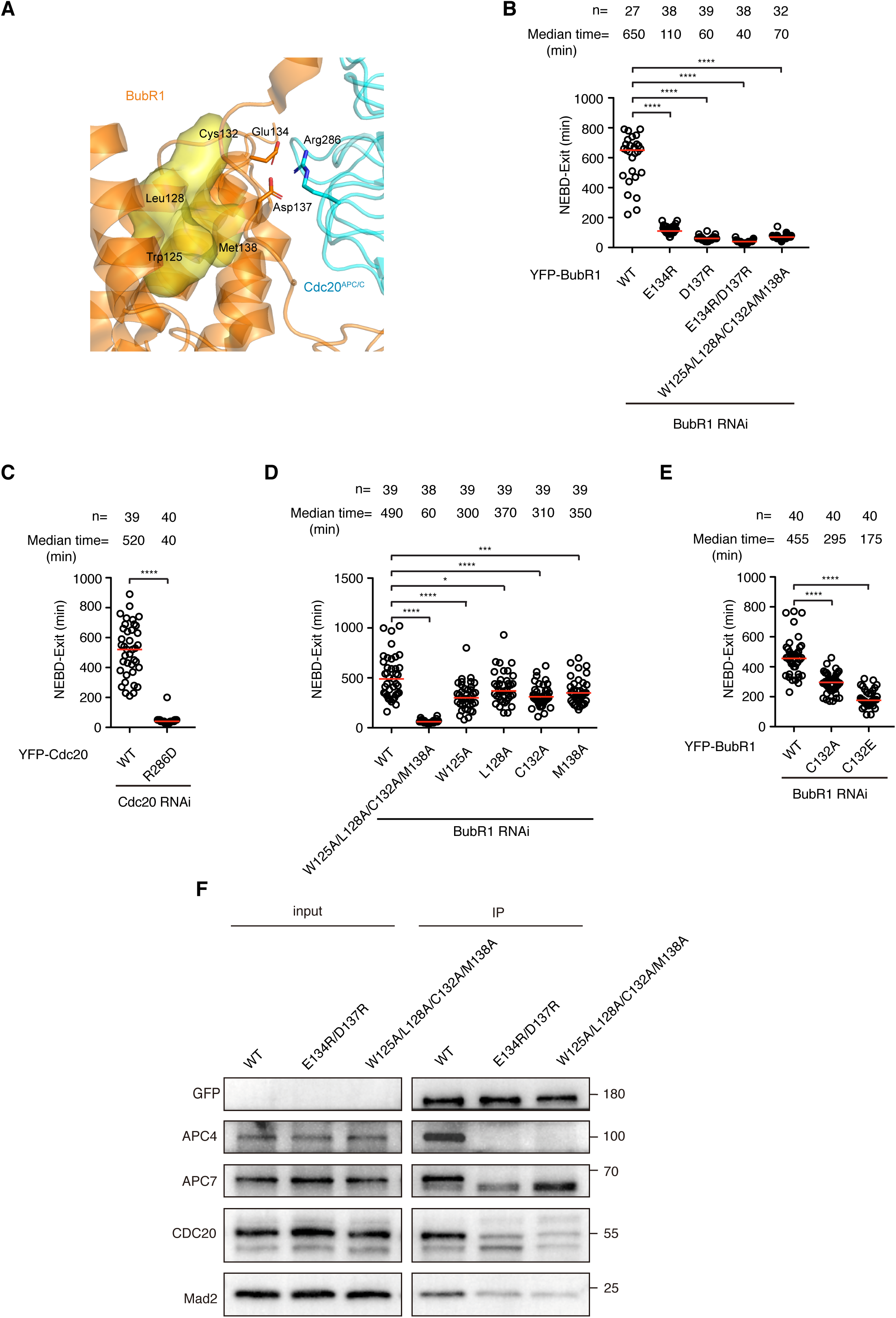
The BubR1 TPR-Cdc20^APC/C^ interaction is critical for SAC signaling. **A** Structural analysis of BubR1 TPR-Cdc20^APC/C^ interaction from PDB: 6TLJ. **B-E** Plots showing the time from NEBD to mitotic exit of the cells transfected with siRNA oligos against BubR1 or Cdc20 and RNAi-resistant constructs expressing YFP-BubR1 or YFP-Cdc20 or corresponding mutants. Low dose of nocodazole (40 ng/ml) was applied into the medium before live cell imaging conducted. Each circle represents the time spent in mitosis of a single cell. Red line indicates the median time. The number of cells analyzed per condition is indicated above (n = X). Mann-Whitney u-test was applied. * means P<0.1; *** means P<0.001; **** means P<0.0001. **F** YFP-BubR1 or YFP-BubR1 E134R/D137R or YFP-BubR1 W125A/L128A/C132A/M138A was expressed in HeLa cells and immunoprecipitated by GFP-trap beads. YFP-BubR1, APC4, APC7, Cdc20 and Mad2 were detected by western blot from the immunoprecipitate.

Notably, this electrostatic interaction lies within a hydrophobic pocket composed of residues previously implicated in binding the KNL1 KI2 motif (Bolanos-Garcia et al., 2011; Krenn et al., 2012) (Fig. 4A, yellow region). This local hydrophobic environment is likely to strengthen the electrostatic interaction between the BubR1 TPR domain and Cdc20. Indeed, a BubR1 mutant with four hydrophobic residues substituted (W125A/L128A/C132A/M138A, termed 4A) reduced checkpoint strength to a level comparable to the E134R/D137R mutant (Fig. 4B). Like the E134R/D137R mutant, this 4A mutant still supported MCC formation, but was defective in APC/C binding (Fig. 4F). Individual alanine substitutions at these hydrophobic residues only modestly impaired SAC signaling (Fig. 4D,E). In contrast, replace one of these residues, Cys132, with a negatively charged glutamate further weakened checkpoint activity, consistent with the role of these residues in maintaining a hydrophobic microenvironment (Fig. 4E). Immunofluorescence analysis confirmed that the W125A/L128A/C132A/M138A mutant exhibited no obvious kinetochore localization defects (Supp. Fig. 1).

### The BubR1 TPR domain binds downstream residues to form a lasso-like structure for capturing Cdc20^APC/C^

Another notable interaction revealed by structural analysis is an intramolecular contact between BubR1 TPR residues and several residues preceding the KEN2 box, specifically, Y141:W295 and E103:R302 (Fig. 5A). This interaction appears to organize a lasso-like structure that encloses or positions nearby all three major Cdc20^APC/C^ interacting motifs: D1(224-232), ABBA1 (272-277), and KEN2 (304-306), along with the newly identified E134/D137 interaction site described above. To validate this interaction, we synthesized biotin-labeled peptides spanning residues Q293 to R302, containing either wild type W295 or a W295A substitution, and performed peptide binding assay with recombinant BubR1 TPR protein. The W295 peptide recruited nearly five times more BubR1 TPR protein than the W295A peptide, indicating that the Y141:W295 interaction is a key driver of the intramolecular binding (Fig. 5B).

**Figure 5.**
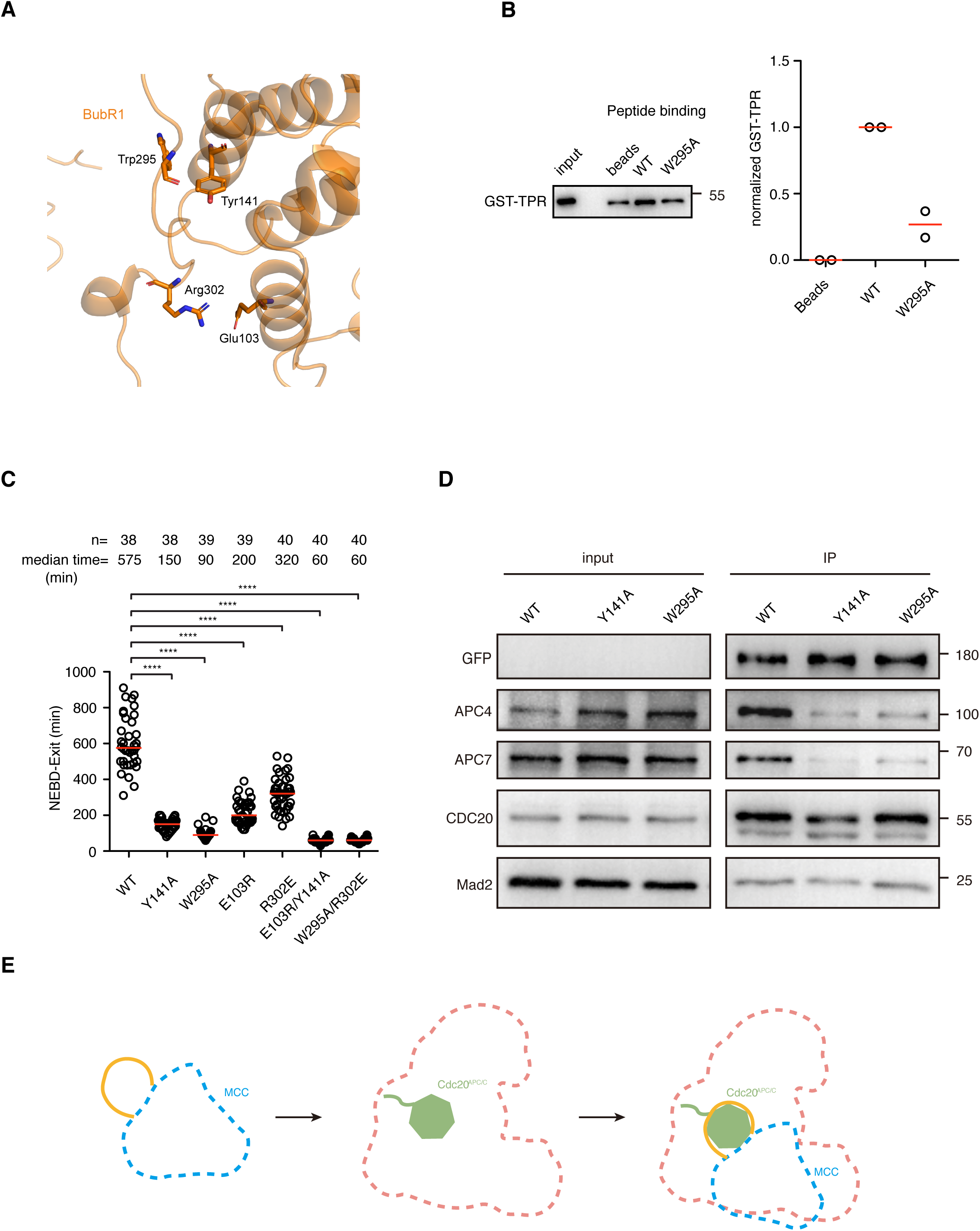
Intramolecular binding between BubR1 TPR and downstream residues may form a lasso structure for Cdc20^APC/C^ capturing. **A** Structural analysis of BubR1 TPR-Mad2 interaction from PDB: 6TLJ. **B** Peptide binding showing a direct binding of the peptide mimicing the short protein sequence in front of BubR1 KEN2 and GST-BubR1 TPR or GST-BubR1 TPR W295A recombinant protein. Left: western blot of the recombinant GST-tagged BubR1 TPR protein bound to the peptides. Right: quantification of the western blot results from two repeats. The line represents the median value of two repeats. The value from beads alone was substracted from the values from the values from the two peptides and set to zero. The value from wild type peptide was set to 1 and the value from peptide W295A was normalized to the wild type value. **C** Plot showing the time from NEBD to mitotic exit of the cells transfected with siRNA oligos against BubR1 and RNAi-resistant constructs expressing YFP-BubR1 or corresponding mutants. Low dose of nocodazole (40 ng/ml) was applied into the medium before live cell imaging conducted. Each circle represents the time spent in mitosis of a single cell. Red line indicates the median time. The number of cells analyzed per condition is indicated above (n = X). Mann-Whitney u-test was applied. **** means P<0.0001. **D** YFP-BubR1 or YFP-BubR1 Y141A or YFP-BubR1 W295A was expressed in HeLa cells and immunoprecipitated by GFP-trap beads. YFP-BubR1, APC4, APC7, Cdc20 and Mad2 were detected by western blot from the immunoprecipitate. **E** Cartoon showing the presence of the lasso structure formed from the intramolecular binding on BubR1 of MCC complex and the full occupation of Cdc20^APC/C^ in APC/C complex.

We next asked whether this intramolecular interaction contributes to SAC signaling. Functional analysis revealed that individual mutations of the four residues impaired checkpoint strength to varying degrees, whereas double mutations (E103R/Y141A or W295A/R302E) nearly abolished SAC activity (Fig. 5C), despite maintaining proper kinetochore localization (Supp. Fig. 1). Immunoprecipitation assay confirmed that MCC assembly remained efficient in the Y141A or W295A single mutants, but their binding to APC/C was severely impaired (Fig. 5D).

These results support a model in which a lasso-like structure in BubR1 molecule facilitates efficient capture of Cdc20^APC/C^ by presenting multiple interaction sites in a spatially coordinated manner, rather than as a linear sequence (Fig. 5E). Further studies are needed to fully validate this model. Within the proposed lasso structure, we also identified a potential interaction between BubR1 residues NNSR (268-271) and Apc10, located upstream of the ABBA1 motif (Supp. Fig. 2). Mutation of NNSR into alanines (4A) caused a mild reduction of the checkpoint signaling and MCC-APC/C binding, without affecting MCC assembly (Supp. Fig. 2).

In summary, our findings demonstrated that the BubR1 TPR domain plays an essential role in SAC signaling by engaging in multiple interactions with APC/C^Cdc20^ and MCC components including BubR1 itself.

## Discussion

The molecular mechanism of SAC signaling has advanced considerably over the last three decades, yet several fundamental questions remain unresolved. In this study, we demonstrated that the BubR1 TPR domain plays an essential role in SAC by engaging in a network of interactions with components of the MCC and APC/C, including Mad2, Cdc20^APC/C^, Apc2 and intramolecularly with BubR1 itself. These interactions collectively enhance MCC formation, stabilize MCC-APC/C binding and efficiently block Cdc20^APC/C^ from its substrates, thereby enabling a rapid SAC activation in response to defective microtubule-kinetochore attachment.

Although a direct interaction between BubR1^1-371^ with Mad2 was previously reported (Tipton et al., 2011), the precise structural determinants and functional relevance of this interaction remained unclear. Guided by the resolved cryo-EM MCC-APC/C structure, we have now precisely mapped the BubR1-Mad2 binding interface and established its functional importance in SAC signaling. Disruption of this interaction markedly enhanced the kinetochore localization of both Mad2 and Cdc20, indicating that the BubR1-Mad2 interaction occurs on kinetochores *in vivo*. Notably, the Mad2 residues involved in binding BubR1 locate at the Mad2 dimeric interface and are required for Mad2 dimerization (Mapelli et al., 2007). This suggests that BubR1 may compete with the Mad2-Mad1 complex to facilitate MCC assembly or release. Further studies will be needed to validate this model.

Another interesting finding of this study is the identification of a hydrophobic pocket within the BubR1 TPR domain which enhances the electrostatic interaction between BubR1 residues E134/D137 and Cdc20^APC/C^ R286. Although the residues such as W125, L128, C132, and M138, which form this pocket, were previously implicated in the BubR1-KNL1 KI2 interaction, their contribution to BubR1 kinetochore appears limited. This was supported by earlier studies mutating KNL1 KI2 motif and confirmed here through direct mutation of BubR1. Indeed, mutation of the KI2 motif in full length KNL1 had minimal effect on the SAC signaling, whereas simultaneous mutation of all four corresponding residues in BubR1 almost fully abolished the SAC activity. Furthermore, introducing a negatively charged amino acid at these sites disrupted checkpoint function more severely than a neutral substitution. Together, these results indicate that the hydrophobic pocket facilitates SAC signaling primarily by strengthening the electrostatic interaction between the BubR1 TPR domain and Cdc20^APC/C^.

A particularly intriguing finding of this study is the identification of a lasso-like structure formed by an intramolecular interaction between BubR1 TPR domain and downstream residues W295 and R302, locate near the KEN2 motif. Disruption of this interaction severely compromises the checkpoint signaling by impairing MCC-APC/C binding. Notably, AlphaFold-predicted structure of BubR1 does not reveal this lasso conformation in the isolated protein, suggesting that it may form during MCC assembly. Previous phylogenetic analysis has shown that W295 and R302 are highly conserved down to the yeast (Tromer et al., 2016). However, the available yeast MCC structure (PDB:4AEZ) lacks the region preceeding KEN2 motif (Chao et al., 2012), preventing direct structural confirmation of the lasso in MCC context. Should this structure indeed form within the MCC, it could enhance the occupancy of the Cdc20^APC/C^ molecule by spatially coordinating four distinct binding sites including D1 box, ABBA1 motif, KEN2 box and E134/D137 electrostatic interface for the binding more efficiently. Although our immunoprecipitation assay clearly showed the defect at pulling down APC/C^Cdc20^ by lasso-defective mutant, future molecular dynamic simulation could provide deeper mechanistic insight into this proposed structural mechanism.

## Materials and methods

### Cell culture and transfection

HeLa cells were cultivated in DMEM medium supplemented with 10% FBS and Pen/Strep. Cell cycle synchronization was performed by double thymidine block. siRNA oligo with RNAi-resistant constructs were transfected into cells inbetween the two thymidine block using Lipofectamine 2000 (Thermo Fisher Scientific). For live cell imaging, the cells released from the second thymidine block were recorded by microscopy 6 hr later. For immunofluorescence assay, cells released from the second thymidine block were treated with RO3306 (5 μM, Selleck) for 12 hr. Afterwards, the cells were released into nocodazole (200 ng/ml, Selleck)-containing DMEM mediuim for 45 min before fixation. RNAi oligos 5’CGGAAGACCUGCCGUUACAtt3’ (Cdc20), 5’GAUGGUGAAUUGUGGAAUAtt3’ (BubR1) and 5’CGUACGCGGAAUACUUCGAtt3’ (luciferase) were synthesized from Genepharma.

### Molecular cloning

The expression plasmids used in this study were generated by standard cloning method and mutagenesis was conducted by mutation PCR. Briefly, wild type BubR1 or Cdc20 was cloned into pcDNA5/FRT/TO N-YFP vector by KpnI and NotI. Gene amplification or mutation PCR was performed with KOD DNA polymerase (Toyobo). All the restriction enzymes and T4 DNA ligase were purchased from Thermo Fisher Scientific.

### Live cell imaging

HeLa cells were seeded in 6-well plate and synchronized with double thymidine block. RNA interference and plasmid transfection was performed as described above. 900 ng of RNAi-resistant YFP-BubR1 or 450 ng of YFP-Cdc20 construct were co-transfected together with 50 nM of respective RNAi oligos in each well. 24 hr later, the cells were re-seeded in 8-well chamber slide (Ibidi) during the second thymidine block. Live cell imaging was performed 6 hr after the cells released from the second thymidine block. The culture medium was replaced by Leibovitz’s L-15 medium (Thermo Fisher Scientific) supplemented with 10% FBS before the slide was mounted onto microscopy. For SAC assays, nocodazole (40 ng/ml, Selleck) was added into the L15 medium. DIC and YFP signals were collected every 10 min for a total of 18 hr by Nikon A1HD25 imaging system (Nikon). NIS-Elements AR Analysis (Nikon) was used for data analysis. At least two repeats for each assay were performed with more than 30 cells in each condition quantified.

### Immunofluorescence

Cells growing on coverslips in 6-well plate were treated as described above. 4% paraformaldehyde in PHEM buffer (60 mM PIPES, 25 mM HEPES, pH 6.9, 10 mM EGTA and 4 mM MgSO_4_) was prepared freshly and applied to the cells for 20 min at room temperature. Afterwards, 0.5% Triton X-100 in PHEM was used to permeabilize cells for 10 min at room temperature followed by cell staining with respective antibodies. The antibodies used in this study include GFP booster Alexa Fluor 488 (Proteintech, gb2AF488-50, 1:500), Cdc20 (Santa Cruz, sc-13162, 1:200), Mad2 (Bethyl, A300-301A, 1:400) and CENP-C (MBL, PD030, 1:800). Fluorescent secondary antibodies were used for the immunofluorescence imaging (Invitrogen, 1:1000) except GFP booster Alexa Fluor 488 was used for YFP detection. Z-stacks with 200 nm interval were taken by DMi8 fluorescent microscopy (Leica) using a 100 × oil objective followed by deconvolution by Las X (Leica). Signal quantification was performed by drawing a circle around each kinetochore marked by CENP-C staining. The three continuous peak values within the circle on the interested channel were averaged and subtracted of the background values from a neighboring circle. Two repeats were performed for each immunofluorescence assay with at least 180 kinetochores from 12 cells in one repeat quantified.

### Immunoprecipitation and western blot

HeLa cells were treated similarly as above for synchronization and transfection. The cells were further treated with nocodazole (200 ng/ml) for an additional 12 hr after released from the second thymidine arrest. Mitotic cells were collected by shake off and lysed in lysis buffer (10 mM Tris pH 7.5, 150 mM NaCl, 0.5 mM EDTA and 0.5% NP40). After centrifugation at 16,000 × g for 15 min at 4 °C, the supernatant was incubated with GFP-Trap beads (Proteintech) and shaken at 1100 × rpm for 2 hr at 4 °C on thermomixer (Eppendorf). After three washes, the bound protein was eluted by 2 × SDS sample buffer and separated by SDS-PAGE. Target proteins were detected by western blot. Antibodies used in this study include YFP antibody (homemade in JN lab), Cdc20 (Santa Cruz, sc-13162), Mad2 (Bethyl, A300-301A), Apc7 (Santa Cruz, sc-365649), Apc4 (Santa Cruz, sc-514895) and GST (Proteintech, 60001-2-lg). HRP-tagged secondary antibody was used for chemiluminescence detection (Proteintech, SA00001-1, SA00001-2).

### Recombinant protein production and protein binding assay

Human Mad2 cDNA (L13A) was cloned into the pCold I vector. Plasmids were transformed into *E. coli* BL21(DE3) cells. After overnight growth of single colony in LB medium (100 μg/mL Ampicillin) at 37°C, bacterial culture was diluted at 1:100 into fresh LB and incubated at 37°C, 200 × rpm until OD₆₀₀ reached 0.6. Protein expression was induced by 0.1 mM IPTG at 16°C for 18 hr. Cells were harvested by centrifugation and resuspended in lysis buffer (20 mM Tris-HCl, pH 8.0, 300 mM NaCl, 5% glycerol). After sonication on ice, the cell lysate was centrifuged at 12,000 × rpm for 30 min at 4°C. The supernatant was incubated with Ni-NTA resin (MCE, HY-K0210) at 4°C for 4 hr with rotation. Mad2 L13A protein was eluted with buffer containing 250 mM imidazole.

The cDNA of human BubR1 TPR (residues 1–230) and BubR1 TPR W22A/E23A were cloned into the pGEX-4T-1 vector. Plasmids were transformed into *E. coli* BL21(DE3). Protein induction and purification procedures were the same as described above except glutathione Sepharose 4B resin (Cytiva, 17075601) was used. After incubation with cell lysate, the resin was washed with PBS and the bound protein was eluted with the elution buffer (50 mM Tris-HCl, pH 8.8, 20 mM reduced glutathione, 300 mM NaCl, and 0.1% Triton X-100). Protein purity was analyzed by SDS-PAGE and the recombinant protein was stored at −80°C till use.

For the binding assay, 15 μl of glutathione Sepharose 4B beads were washed three times with PBS and resuspended in 500 μl of PBS. Afterwards, 1 μg of GST-tagged BubR1 TPR recombinant protein was incubated with the beads suspension overnight at 4°C with rotation. After incubation, 1 μg of purified His-Mad2 L13A protein was added and the mixture was incubated for 4 hr at 4°C with rotation. Beads were washed 4 times with PBS and the bound proteins were eluted in 40 μl of 2× SDS loading buffer by heating at 95°C for 5 min. Eluted samples were analyzed by western blotting with GST antibody (Proteintech, 66001-2-lg) and His antibody (Proteintech, 66005-1-lg).

### Peptide pull-down assay

Biotinylated peptides (BubR1 WT: QPWIAPPMPRW; BubR1 W295A: QPAIAPPMPRW) were synthesized by Sangon and dissolved in DMSO at 4 mg/ml. 30 μl of streptavidin resin (Thermo Scientific) was washed three times by PBST (PBS with 0.1% Tween-20). The resin was then incubated with 5 μl of the biotinylated peptide at room temperature for 1 hr with rotation. After incubation, the resin was washed twice with PBST to remove unbound peptide. 3 μg of recombinant BubR1 TPR protein was diluted in 500 μL of lysis buffer (50 mM Tris-HCl (pH 7.5), 150 mM NaCl, 0.3% Triton X-100, supplemented with protease and phosphatase inhibitors) and added to the peptide-loaded resin. The mixture was incubated at 24 °C for 1 hr with rotation. Afterwards, the resin was washed three times with PBST. The bound protein was eluted by adding 50 μL of 2× SDS loading buffer onto the resin and heating at 95 °C for 5 min. Eluted samples were analyzed by western blot with GST antibody (Proteintech, 66001-2-lg) and His antibody (Proteintech, 66005-1-lg).

### Statistics and Reproducibility

Mann-Whitney u-test was applied for all the statistic analysis for live imaging results, immunofluorescence quantification and peptide intensity which have been repeated at least twice. ns means not significant; * means P<0.1; ** means P<0.01; *** means P<0.001; **** means P<0.0001.

---

**Supplementary Figure 1.**
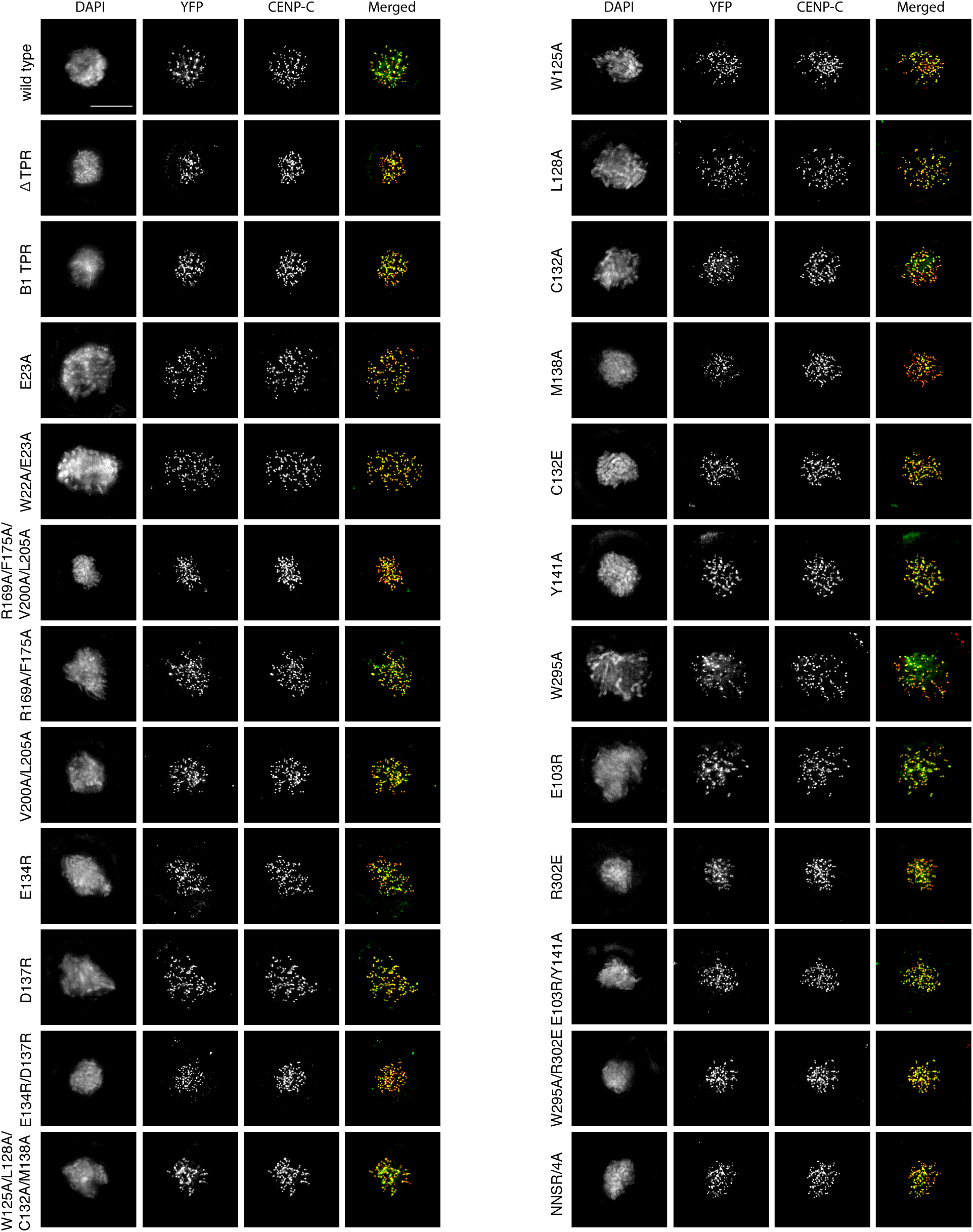
Kinetochore localization of BubR1 mutants. Immunofluorescence images of HeLa cells expressing wild type or mutant YFP-BubR1 with endogenous BubR1 depleted by RNAi. Merged images showing YFP-BubR1 in green and CENP-C in red.

**Supplementary Figure 2.**
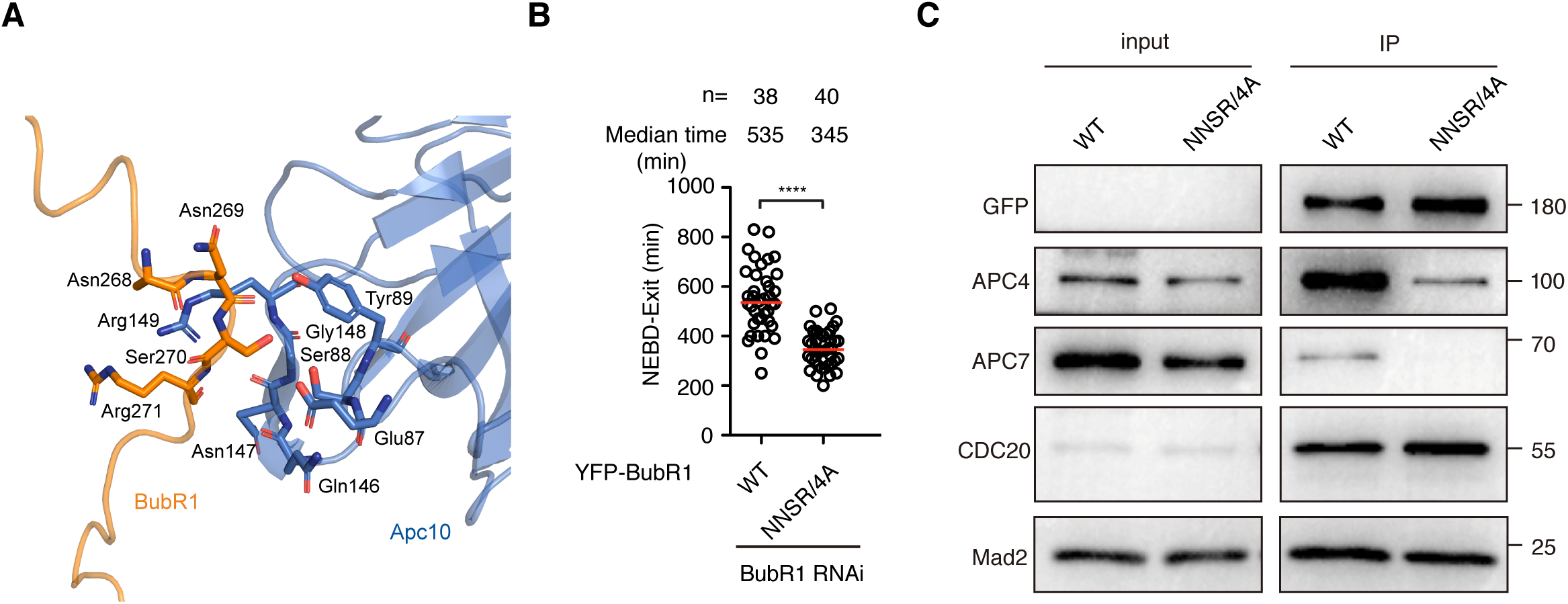
BubR1 TPR-Apc10 interaction is required for full activation of SAC. **A** Structural analysis of BubR1 TPR-Apc10 interaction from PDB: 6TLJ. **B** Plot showing the time from NEBD to mitotic exit of the cells transfected with siRNA oligos against BubR1 and RNAi-resistant constructs expressing YFP-BubR1 or YFP-BubR1 NNSR/4A. Low dose of nocodazole (40 ng/ml) was applied into the medium before live cell imaging conducted. Each circle represents the time spent in mitosis of a single cell. Red line indicates the median time. The number of cells analyzed per condition is indicated above (n = X). Mann-Whitney u-test was applied. **** means P<0.0001. **C** YFP-BubR1 or YFP-BubR1 NNSR/4A was expressed in HeLa cells and immunoprecipitated by GFP-trap beads. YFP-BubR1, APC4, APC7, Cdc20 and Mad2 were detected by western blot from the immunoprecipitate.

## Acknowledgement

We thank Dr. Claudio Alfieri at the Institute of Cancer Research, London and Prof. Jakob Nilsson at the Danish Cancer Society for constructive advices on the project. GZ is supported by the National Natural Science Foundation of China (31970666), Taishan Scholar Project (tsqn201812054) from Shandong, China and Shandong Provincial Natural Science Foundation (ZR2025SM287).

## References

Musacchio, A. The molecular biology of spindle assembly checkpoint signaling dynamics. Curr. Biol. 25, R1002–R1018 (2015).

Saurin, A.T. Kinase and phosphatase cross-talk at the kinetochore. Front. Cell Dev. Biol. 6, 62 (2018).

Ji, Z., Gao, H., Jia, L., Li, B. & Yu, H. A sequential multi-target Mps1 phosphorylation cascade promotes spindle checkpoint signaling. eLife 6, e22513 (2017).

Zhang, G. et al. Bub1 positions Mad1 close to KNL1 MELT repeats to promote checkpoint signalling. Nat. Commun. 8, 15822 (2017).

Faesen, A.C. et al. Basis of catalytic assembly of the mitotic checkpoint complex. Nature 542, 498–502 (2017).

Lara-Gonzalez, P., Kim, T., Oegema, K., Corbett, K. & Desai, A. A tripartite mechanism catalyzes Mad2-Cdc20 assembly at unattached kinetochores. *Science* **371**, 64-67 (2021). Piano, V. et al. CDC20 assists its catalytic incorporation in the mitotic checkpoint complex. Science 371, 67–71 (2021).

Fischer, E.S. et al. Molecular mechanism of Mad1 kinetochore targeting by phosphorylated Bub1. EMBO Rep. 22, e52242 (2021).

Fischer, E.S. et al. Juxtaposition of Bub1 and Cdc20 on phosphorylated Mad1 during catalytic mitotic checkpoint complex assembly. Nat. Commun. 13, 6381 (2022).

Chen, C. et al. The structural flexibility of MAD1 facilitates the assembly of the mitotic checkpoint complex. Nat. Commun. 14, 1529 (2023).

Sethi, S. et al. Interplay of kinetochores and catalysts drives rapid assembly of the mitotic checkpoint complex. Nat. Commun. 16, 4823 (2025).

Yu, C.W.H. et al. Molecular mechanism of Mad2 conformational conversion promoted by the Mad2-interaction motif of Cdc20. Protein Sci. 34, e70099 (2025).

Kulukian, A., Han, J.S., Cleveland, D.W. Unattached kinetochores catalyze production of an anaphase inhibitor that requires a Mad2 template to prime Cdc20 for BubR1 binding. Dev. Cell 16, 105–117 (2009).

Burton, J.L. & Solomon, M.J. Mad3p, a pseudosubstrate inhibitor of APCCdc20 in the spindle assembly checkpoint. Genes Dev. 21, 655–667 (2007).

King, E.M.J., van der Sar S.J.A. & Hardwick, K.G. Mad3 KEN boxes mediate both Cdc20 and Mad3 turnover, and are critical for the spindle checkpoint. PLoS One 2, e342 (2007).

Sczaniecka, M. et al. The spindle checkpoint functions of Mad3 and Mad2 depend on a Mad3 KEN box-mediated interaction with Cdc20-anaphase-promoting complex (APC/C). J. Biol. Chem. 283, 23039–23047 (2008).

Malureanu, L.A., Jeganathan, K.B., Hamada, M., Wasilewski, L., Davenport, J. & van Deursen, J.M. BubR1 N terminus acts as a soluble inhibitor of cyclin B degradation by APC/C(Cdc20) in interphase. Dev. Cell 16, 118–131 (2009).

Elowe, S., Dulla, K., Uldschmid, A., Li, X., Dou, Z. & Nigg, E.A. Uncoupling of the spindle-checkpoint and chromosome-congression functions of BubR1. J. Cell Sci. 123, 84–94 (2010).

Lara-Gonzalez, P., Scott, M.I.F., Diez, M., Sen, O. & Taylor, S.S. BubR1 blocks substrate recruitment to the APC/C in a KEN-box-dependent manner. J. Cell Sci. 124, 4332–4345 (2011).

Lischetti, T., Zhang, G., Sedgwick, G.G., Bolanos-Garcia, V.M. & Nilsson, J. The internal Cdc20 binding site in BubR1 facilitates both spindle assembly checkpoint signalling and silencing. Nat. Commun. 5, 5563 (2014).

Di Fiore, B. et al. The ABBA motif binds APC/C activators and is shared by APC/C substrates and regulators. Dev. Cell 32, 358–372 (2015).

Diaz-Martinez, L.A. et al. The Cdc20-binding Phe box of the spindle checkpoint protein BubR1 maintains the mitotic checkpoint complex during mitosis. J. Biol. Chem. 290, 2431–2443 (2015).

Izawa, D. & Pines, J. The mitotic checkpoint complex binds a second CDC20 to inhibit active APC/C. Nature 517, 631–634 (2015).

Di Fiore, B., Wurzenberger, C., Davey, N.E. & Pines, J. The mitotic checkpoint complex requires an evolutionary conserved cassette to bind and inhibit active APC/C. Mol. Cell 64, 1144–1153 (2016).

Alfieri, C. et al. Molecular basis of APC/C regulation by the spindle assembly checkpoint. Nature 536, 431–436 (2016).

Sewart, K. & Hauf, S. Different functionality of Cdc20 binding sites within the mitotic checkpoint complex. Curr. Biol. 27, 1213–1220 (2017).

Taylor, S.S., Ha, E. & McKeon F. The human homologue of Bub3 is required for kinetochore localization of Bub1 and a Mad3/Bub1-related protein kinase. J. Cell Biol. 142, 1–11 (1998).

Wang, X. et al. The mitotic checkpoint protein hBUB3 and the mRNA export factor hRAE1 interact with GLE2p-binding sequence (GLEBS)-containing proteins. J. Biol. Chem. 276, 26559–26567 (2001).

Larsen, N.A., Al-Bassam, J., Wei, R.R. & Harrison, S.C. Structural analysis of Bub3 interactions in the mitotic spindle checkpoint. Proc. Natl. Acad. Sci. U. S. A. 104, 1201–1206 (2007).

Zhang, G., Lopez-Mendez, B., Sedgwick, G.G. & Nilsson, J. Two functionally distinct kinetochore pools of BubR1 ensure accurate chromosome segregation. Nat. Commun. 7, 12256 (2016).

Suijkerbuijk, S.J.E., Vleugel, M., Teixeira, A., and Kops, G.J.P.L. Integration of kinase and phosphatase activities by BubR1 ensures formation of stable kinetochore-microtubule attachment. Dev. Cell 23, 745–755 (2012).

Kruse, T. et al. Direct binding between BubR1 and B56-PP2A phosphatase complexes regulate mitotic progression. J. Cell Sci. 126, 1086–1092 (2013).

Xu, P., Raetz, E.A., Kitagawa, M., Virshup, D.M. & Lee, S.H. BUBR1 recruits PP2A via the B56 family of targeting subunits to promote chromosome congression. Biol. Open 2, 479–486 (2013).

Kiyomitsu, T., Obuse, C. & Yanagida, M. Human Blinkin/AF15q14 is required for chromosome alignment and the mitotic checkpoint through direct interaction with Bub1 and BubR1. Dev. Cell 13, 663–676 (2007).

Kiyomitsu, T., Murakami, H. & Yanagida, M. Protein interaction domain mapping of human kinetochore protein Blinkin reveals a consensus motif for binding of spindle assembly checkpoint proteins Bub1 and BubR1. Mol. Cell Biol. 31, 998–1011 (2011).

Bolanos-Garcia, V.M. et al. Structure of a Blinkin-BUBR1 complex reveals an interaction crucial for kinetochore-mitotic checkpoint regulation via an unanticipated binding site. Structure 19, 1691–1700 (2011).

Krenn, V., Wehenkel, A., Li, X., Santaguida, S. & Musacchio, A. Structural analysis reveals features of the spindle checkpoint kinase Bub1-kinetochore subunit Knl1 interaction. J. Cell Biol. 196, 451–467 (2012).

London, N., Ceto, S., Ranish, J.A. & Biggins, S. Phosphoregulation of Spc105 by Mps1 and PP1 regulates Bub1 localization to kinetochores. Curr. Biol. 22, 900–906 (2012).

Shepperd, L.A. et al. Phosphodependent recruitment of Bub1 and Bub3 to Spc7/KNL1 by Mph1 kinase maintains the spindle checkpoint. Curr. Biol. 22, 891–899 (2012).

Yamagishi, Y., Yang, C.H., Tanno, Y. & Watanabe, Y. MPS1/Mph1 phosphorylates the kinetochore protein KNL1/Spc7 to recruit SAC components. Nat. Cell Biol. 14, 746–752 (2012).

Vleugel, M. et al. Arrayed BUB recruitment modules in the kinetochore scaffold KNL1 promote accurate chromosome segregation. J Cell Biol. 203, 943–955 (2013).

Krenn, V., Overlack, K., Primorac, I., van Gerwen, S. & Musacchio, A. KI motifs of human Knl1 enhance assembly of comprehensive spindle checkpoint complexes around MELT repeats. Curr. Biol. 24, 29–39 (2014).

Tipton, A.R. et al. BUBR1 and closed MAD2 (C-MAD2) interact directly to assemble a functional mitotic checkpoint complex. J. Biol. Chem. 286, 21173–21179 (2011).

Tromer, E., Bade, D., Snel, B. & Kops, G.J.P.L. Phylogenomics-guided discovery of a novel conserved cassette of short linear motifs in BubR1 essential for the spindle checkpoint. Open Biol. 6, 160315 (2016).

Fujita, H., et al. Premature aging syndrome showing random chromosome number instabilities with CDC20 mutation. Aging Cell 19, e13251 (2020).

Tian, W., Li, B., Warrington, R., Tomchick, D.R., Yu, H. & Luo, X. Structural analysis of human Cdc20 supports multisite degron recognition by APC/C. Proc. Natl. Acad. Sci. U.S.A. 109, 18419–18424 (2012).

Alfieri, C., Tischer, T. & Barford, D. A unique binding mode of Nek2A to the APC/C allows its ubiquitination during prometaphase. EMBO Rep. 21, e49831 (2020).

Song, C. et al. Self-priming of Plk1 binding to BubR1 ensures accurate mitotic progression. *Commun*. Biol. 7, 1473 (2024).

Mapelli, M., Massimiliano, L., Santaguida, S. & Musacchio, A. The Mad2 conformational dimer: structure and implications for the spindle assembly checkpoint. Cell 131, 730–743 (2007).

Chao, W.C.H., Kulkarni, K., Zhang, Z., Kong, E.H. & Barford, D. Structure of the mitotic checkpoint complex. Nature 484, 208–213 (2012).

